# Differential value processing in the rostral and caudal subthalamic nucleus during flexible and habitual decision-making in primates: Implications for cognitive dysfunction in Parkinson’s disease

**DOI:** 10.64898/2026.10.19.760032

**Authors:** Keonwoo Lee, Myong Lee, Haiyan Jiang, Hyoung F. Kim

## Abstract

The subthalamic nucleus (STN) is a major target region of deep brain stimulation (DBS) for Parkinson’s disease (PD), effectively alleviating motor symptoms. However, STN-DBS can also induce cognitive impairments, suggesting that the STN contributes to cognitive functions beyond simple motor control. Furthermore, anatomical organization supports functional segregation of cognitive processes, as basal ganglia subregions encoding flexible and stable values topographically project to distinct STN subregions. Yet, whether the STN contributes to cognitive functions and, if so, how subregions of the STN differentially process cognitive flexibility and habitual stability remains unknown. Here, we reveal a rostral–caudal functional organization of the STN that differentially supports flexible value updating and long-term value memory, enabling adaptive and habitual decision-making, respectively. Neurons in the rostral STN (rSTN) preferentially encode flexibly updated values, whereas neurons in the caudal STN (cSTN) preferentially encode stable long-term value memories. Notably, inactivation of the STN along the rostral-caudal axis produced dissociable behavioral deficits: rSTN inactivation selectively impaired goal-directed behavior requiring flexible value updating, whereas cSTN inactivation selectively disrupted habitual behavior guided by long-term value memories. Together, these results reveal a rostral-caudal functional segregation of the STN, with the rSTN supporting goal-directed behavior and the cSTN supporting habitual behavior. These findings further provide a framework for understanding how STN dysfunction or STN-DBS in Parkinson’s disease may differentially affect cognitive processes beyond motor control.

## Introduction

Parkinson’s disease (PD) is a systemic neurodegenerative disorder induced by dopamine depletion, causing motor impairments^1–4^. It is widely known that PD symptoms include tremors and rigidity^5–7^, and circuit mechanisms underlying motor deficits have been suggested: Alterations in basal ganglia circuit activity, followed by degeneration of dopaminergic neurons, lead to heightened neuronal activity in the subthalamic nucleus (STN). The hyperactivity in the STN induces imbalanced responses in the thalamic and cortical regions^8–11^. Therefore, STN inhibition with deep brain stimulation (DBS) has been established as an effective therapy for alleviating motor impairments in PD^12,13^. Furthermore, the relationship between anatomical positions and therapeutic effects of STN-DBS has been investigated^14,15^. Accordingly, the STN has been considered a key structure for motor control. Previous studies have demonstrated that the STN plays a role in switching strategies from automatic to controlled movement^16^. In addition, subthalamic neurons exhibited distinct responses during movement initiation and cancellation^17^.

However, Parkinson’s disease is also associated with a range of non-motor symptoms, including mood disorders and cognitive deficits, in addition to its characteristic motor impairments^18,19^. Clinical research has examined impaired habit learning and working memory in PD patients, accompanied by cognitive disorders^20,21^. In contrast to motor aspects of PD, the basal ganglia circuits underlying the non-motor deficits of PD have not been clearly delineated. Moreover, earlier studies also reported that DBS for motor symptoms could worsen non-motor symptoms; PD patients who underwent STN-DBS showed dysfunction in visual learning and value-based decision-making^22,23^. These observations suggest that the STN may contribute to functions beyond motor control. We therefore hypothesized that the STN is involved not only in motor control but also in cognitive behaviors, including value-based decision-making.

The STN receives GABAergic input from the globus pallidus externa (GPe) and, in turn, sends glutamatergic projections to the substantia nigra pars reticulata (SNr)^24,25^. Notably, the GPe and SNr have been implicated in processing long-term value memories that contribute to habitual behavior^26,27^. The STN also receives dopaminergic inputs from the substantia nigra pars compacta (SNc)^28,29^, whose neurons encode reward-prediction error and value-maintaining signals that guide goal-directed and habitual behaviors^30–32^. Beyond the basal ganglia, the STN receives inputs from the cortex via the hyperdirect pathway^33,34^. In particular, cortical regions such as orbitofrontal cortex (OFC) and ventromedial prefrontal cortex (vmPFC) are known to play key roles in value-based decision-making, suggesting that cortical inputs to the STN may also convey value-related information^35–40^. Together, these anatomical connections between the STN and multiple value-processing structures suggest that the STN plays a role in value-based decision-making.

Moreover, accumulating evidence indicates that multiple basal ganglia structures, including the primate striatum, GPe, and SNr, are organized along a common rostral-caudal functional axis^24,27,41–44^. Along the rostral-caudal axis of these structures, rostral regions preferentially support flexible, goal-directed decision-making, whereas caudal regions preferentially support stable, habitual decision-making. Interestingly, anatomical studies in macaques have shown that pallidal inputs to the STN are topographically organized and that neurons in the caudal STN preferentially project to the caudal SNr^45,46^. Together, these findings indicate that the STN is embedded within a broad topographically organized basal ganglia network, raising the possibility of functional specialization within the STN. Understanding the rostral-caudal organization of the STN may provide critical insights into the circuit mechanisms underlying cognitive impairments in Parkinson’s disease and inform the development of subregion-specific therapies.

From these functional and anatomical observations, we hypothesized that the STN processes value information along its rostral-caudal axis, with distinct subregions differentially contributing to flexible goal-directed and stable habitual decision-making. To test this hypothesis, we recorded neuronal activity throughout the STN while monkeys performed tasks requiring either flexible value updating or stable value-guided behavior, and examined the functional properties and anatomical distribution of individual neurons. We further demonstrated the causal contributions of anatomically defined STN subregions through targeted inactivation experiments. Using this approach, we sought to determine how distinct STN subregions support value-based decision-making and whether their functional organization provides a framework for understanding cognitive dysfunction associated with Parkinson’s disease and STN-DBS.

## Results

### Goal-directed and habitual control of value-based decision-making

Two macaque monkeys performed tasks designed to assess eye movements based on flexible and stable values of visual objects^42,43^. Flexible value was examined using an object value reversal task, in which one fractal object was associated with reward (high-valued) and the other with no reward (low-valued) (Fig. 1A). The object–reward contingency was periodically reversed, requiring continuous updating of object values. As the white fixation dot disappeared, monkeys made a saccadic eye movement toward the fractal, and the reward was delivered according to the object-reward contingency. This association was reversed every 20-25 trials, and the monkey had to exhibit a goal-directed saccade guided by a flexibly changing value. We measured the reaction time (RT) after the disappearance of the fixation dot (go cue), and the monkeys showed faster reaction times toward the high-valued objects than low-valued objects (65.1 ms ± 11.7 ms for high-valued objects and 91.6 ms ± 13.2 ms for low-valued objects) (p = 5.32e-33, Wilcoxon rank-sum test) (Fig. 1B). These results indicate that the monkeys flexibly adapted to changes in object values and exhibited goal-directed behavior.

**Figure 1.**
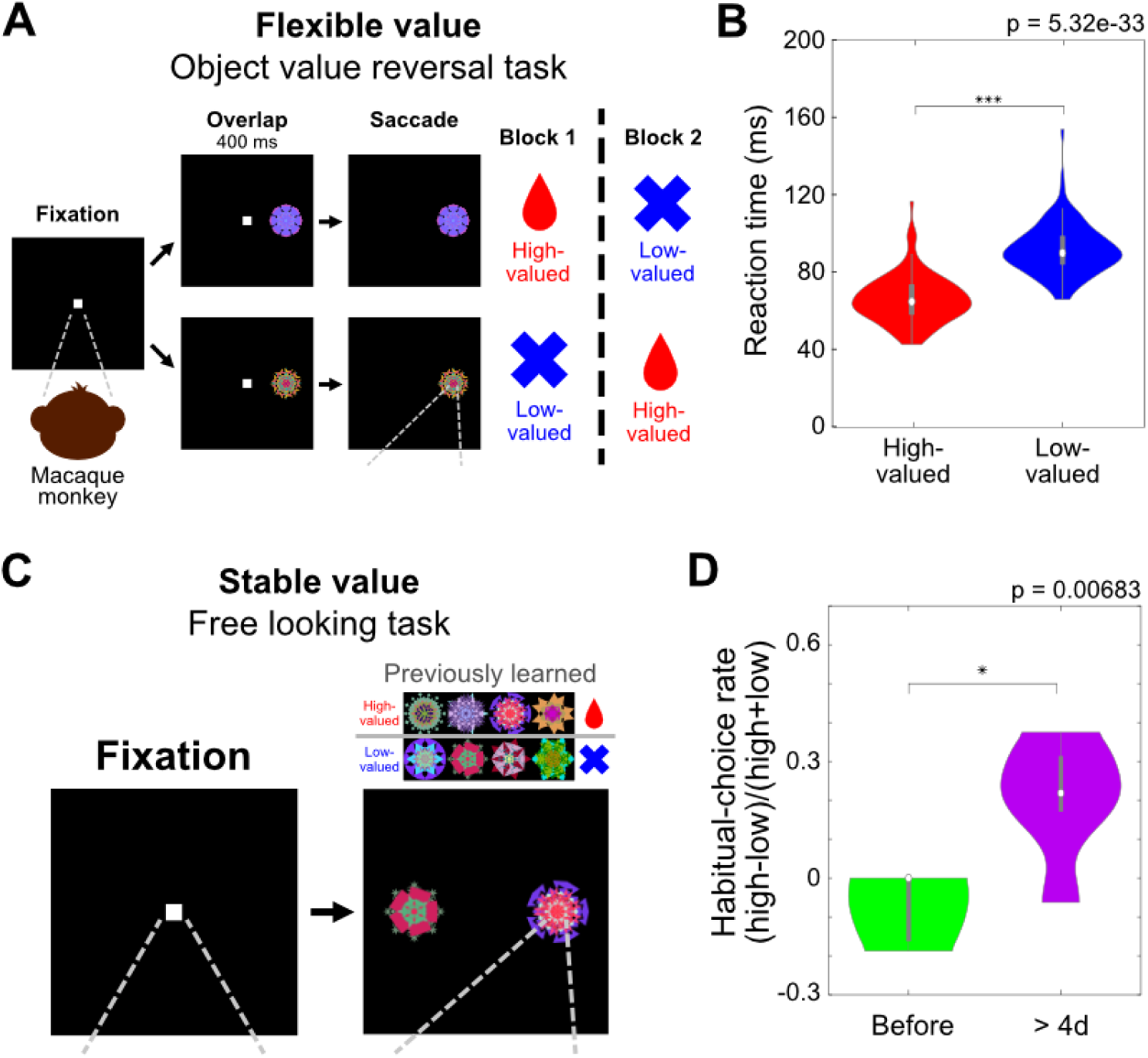
Flexible and stable value procedures and behaviors guided by flexible and stable values. (A) The scheme of the object value reversal task for testing goal-directed behavior. While the monkey was fixating on a central fixation dot, one of the two objects appeared at the left or right position. After 400 ms, as the central fixation dot disappeared, the monkey was forced to make a saccade toward the object. In a block of 20-25 trials, the monkey received a reward after making a saccade to one object, but not the other. (B) The averaged reaction times for the high-valued object and the low-valued object in the object value reversal task. The central white dot in a violin plot indicates the median, and the bottom and the top of the middle gray box correspond to the 25th and 75th percentiles, respectively. (C) The scheme of the free looking task for testing visual habits. Four objects were always associated with a reward (high-valued objects), while the other four objects were always associated with no reward (low-valued objects). In the free looking task, among the 8 previously learned objects, one object from the high-valued objects and one object from the low-valued objects were selected pseudo-randomly from one set and presented simultaneously. The monkey was free to look at these objects (or look elsewhere) for 2 s without any reward feedback. (D) Habitual-choice rate before and after learning. The habitual-choice rate was defined as (high - low) / (high + low), with ‘high’ and ‘low’ referring to the number of saccades toward high-valued and low-valued objects, respectively. The central white dot in a violin plot indicates the median, and the bottom and the top of the middle gray box correspond to the 25th and 75th percentiles, respectively. *p < 0.05, ***p < 0.0005.

To examine habitual behavior, monkeys first learned the values of objects that were consistently associated with reward during a stable value learning task (Fig. S1). This training was repeated for more than four days. After a retention period of at least one day following the final training session, monkeys performed a free looking task to assess habitual behavior guided by long-term stable value memory.

In the free looking task, a high-valued object and a low-valued object from a set of previously long-term learned objects were shown on the left and right sides of the screen. Monkeys were free to view the objects, but no reward was delivered regardless of which object they looked at (Fig. 1C). We compared the proportion of first saccades directed toward high- and low-valued objects before and after more than four days of stable value learning. Monkeys directed their gaze toward high-valued objects more frequently after learning than before learning (habitual choice rate: −0.08 ± 0.07 before learning vs. 0.25 ± 0.10 after more than four days of learning; p = 0.0068, two-tailed paired t-test), despite the absence of reward delivery during the task (Fig. 1D). Together, these behavioral results demonstrate that monkeys flexibly updated object values in the reversal task and formed stable long-term value memories in the stable value learning task, supporting goal-directed and habitual behavior, respectively.

### Flexible and stable value encoding in three example neurons of the STN

In these conditions, we recorded neuronal activity in the STN to examine how flexible and stable values are encoded. In the object value reversal task, we investigated whether STN neurons encode flexible object values that guide goal-directed behavior. To isolate object value information and minimize motor signals related to eye movements, we analyzed neuronal responses to objects during the fixation dot–object overlap phase (Fig. 2A).

**Figure 2.**
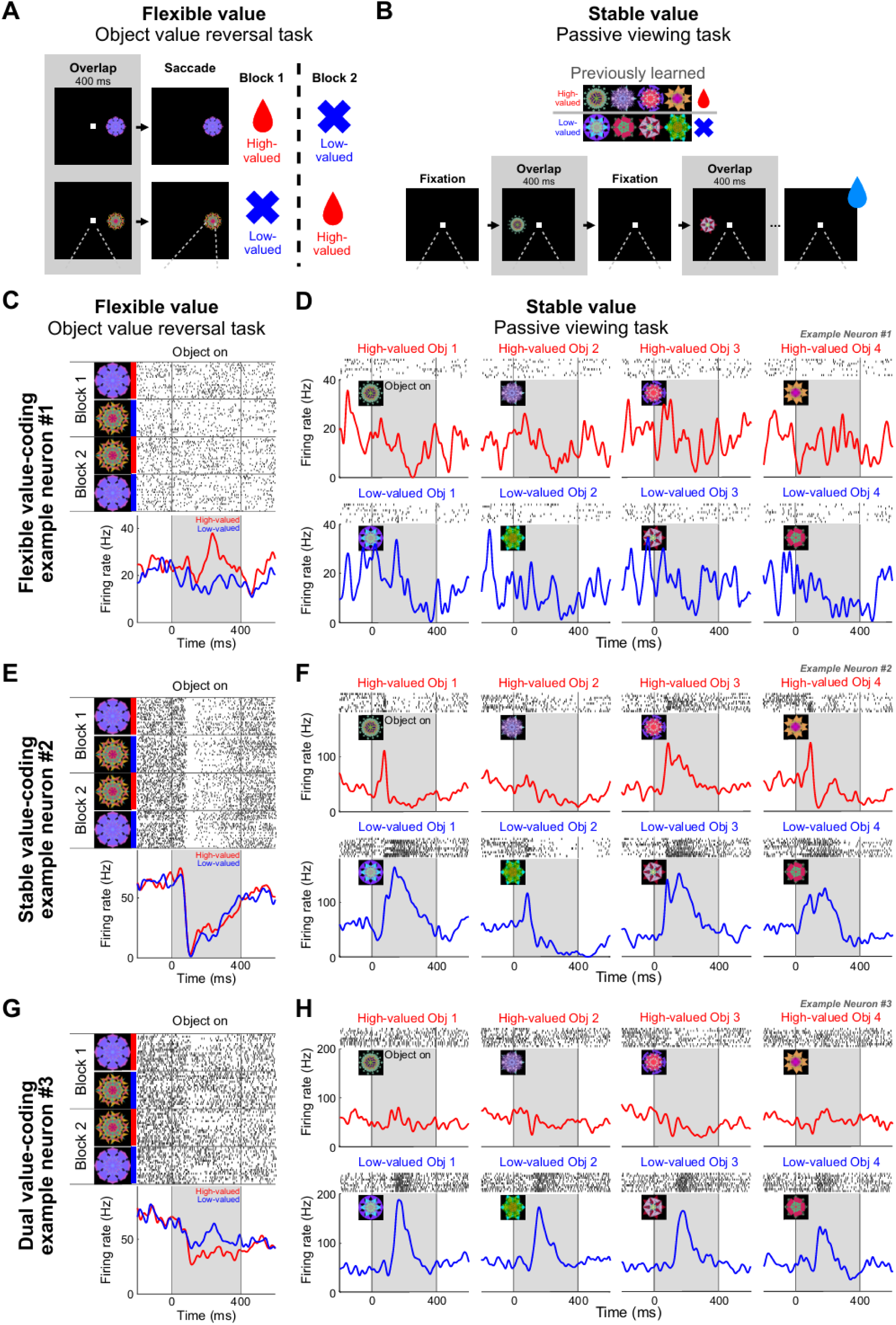
Example neurons in the STN encoding flexible value, stable value, or dual value. (A) Object value reversal task procedure for testing neurons’ responses to objects associated with flexible value (same as in Figure 1A). (B) Passive viewing task procedure for testing neurons’ responses to objects associated with stable value. While the monkey was fixating on a central white dot, two to four fractal objects were chosen pseudo-randomly from a set of 8 previously learned objects and were presented sequentially for 400 ms in the receptive field. The monkey was rewarded 300 ms after the final object disappeared; thus, the reward was not directly associated with any object. (C-H) Representative STN neurons encoding flexible value (C and D), stable value (E and F), and dual value (G and H). Black dots in the raster plots above the peristimulus time histograms indicate neuronal spike responses. The responses of the example STN neurons to fractal objects are associated with flexible value in the object value reversal task (C, E, and G) and stable value in the passive viewing task (D, F, and H), and are aligned to the object onset. The red lines indicate the responses to the high-valued objects, and the blue lines indicate the responses to the low-valued objects.

To assess the encoding of stable value memories in STN neurons, we used a passive viewing task as previously described^32,42,43,47^. In this task, no associative reward was provided, preventing value updating and allowing automatic retrieval of stable object values. While the monkey fixated on a white fixation dot, two to four previously learned objects were sequentially presented within the receptive field (Fig. 2B). We analyzed neuronal responses to objects during the fixation dot-object overlap phase.

Figures 2C and 2D show an example neuron selectively encoding flexible value but not stable value. This neuron showed significantly greater responses to high-valued than to low-valued objects in the object value reversal task (p = 1.1e-05, Wilcoxon rank-sum test), indicating flexible value encoding (Fig. 2C). In contrast, its responses to objects with long-term learned values were not significantly modulated by value in the passive viewing task (p = 0.8725, Wilcoxon rank-sum test) (Fig. 2D). Because this neuron encoded flexible value but not stable value, it was classified as a flexible value-coding neuron.

An example neuron selectively encoding stable value is shown in Figure 2E and 2F. In the object value reversal task, regardless of object value, the neuronal activity decreased upon object presentation. This neuron did not show a significant difference in firing between high-valued and low-valued objects (p = 0.8148, Wilcoxon rank-sum test) (Fig. 2E). In the passive viewing task for testing stable value, this same neuron showed higher and more sustained activity during the presentation of low-valued objects than during the presentation of high-valued objects (p = 1.17e-05, Wilcoxon rank-sum test) (Fig. 2F). Therefore, this neuron was classified as a stable value-coding neuron.

We identified another type of value-coding neuron in the STN that encoded both flexible and stable values. In the object value reversal task, neuronal activity decreased during object presentation in the overlap phase, with significantly stronger inhibition for high-valued than for low-valued objects (p = 0.0031, Wilcoxon rank-sum test) (Fig. 2G). Interestingly, in the passive viewing task, this neuron exhibited significantly stronger responses to stable low-valued than to stable high-valued objects (p = 2.66e-15, Wilcoxon rank-sum test) (Fig. 2H). Because this neuron encoded both flexible and stable values, it was classified as a dual value-coding neuron. Overall, we identified three types of value-coding neurons in the STN: flexible value-coding neurons, stable value-coding neurons, and dual value-coding neurons.

### Population responses support flexible and stable value processing in the STN

We identified 180 task-related neurons that exhibited visual, value-related, or reward-related responses. 46.1% (83/180) of STN neurons showed reward-related or visual responses to the object itself without encoding object value. The other 53.9% (97/180) neurons were value-coding neurons. We classified these value-coding neurons into 3 types based on which value types were encoded, as described in Figure 2: flexible value-coding neurons (48%, n = 47), which exclusively encode flexible value; stable value-coding neurons (25%, n = 24), which selectively encode stable value only; and dual value-coding neurons (27%, n = 26) which encode both flexible and stable values (Fig. 3A).

**Figure 3.**
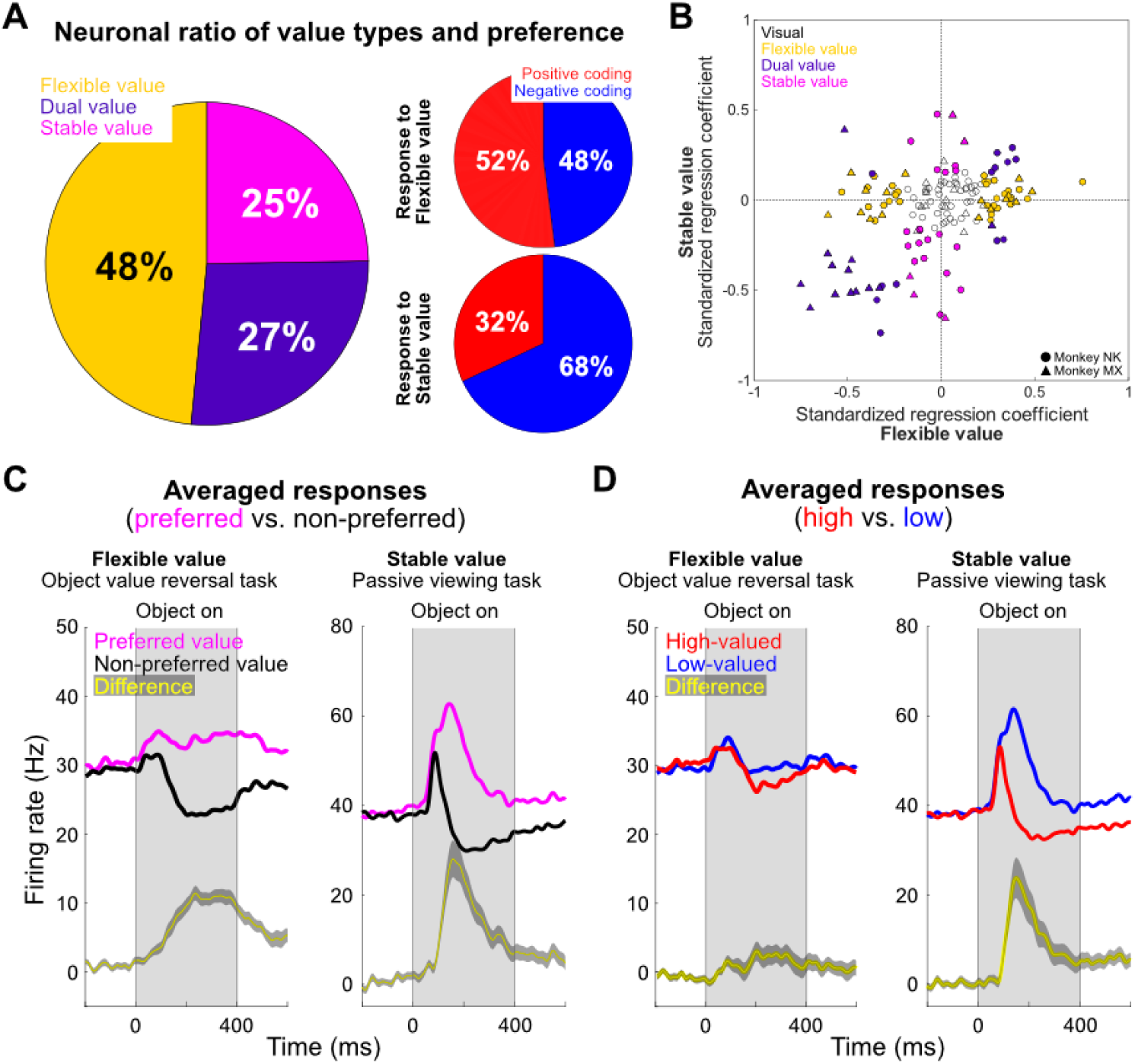
Averaged responses of flexible and stable value in the STN. (A) Pie chart of the recorded STN neurons (n = 180) categorized by value-coding type (left panel). The yellow portion represents flexible value-coding neurons, the purple portion represents dual value-coding neurons, and the magenta portion represents stable value-coding neurons. Pie charts of value-coding neurons processing flexible value (n = 73, upper right) and stable value (n = 50, lower right) categorized by value preference type. The red portion represents the positive coding type and the blue portion represents the negative coding type, which show stronger responses toward high-valued and low-valued objects, respectively. (B) Scatterplot of standardized regression coefficients for the stable value (ordinate) against those for the flexible value (abscissa) in the STN. (C and D) The averaged response of value-coding neurons in the STN (n = 97) during the object value reversal task (left) and the passive viewing task (right). Neuronal responses are aligned to the object onset and averaged for the neurons’ preferred (magenta) and non-preferred (black) values (C), and high-valued (red) and low-valued (blue) objects (D). The yellow line indicates the difference between the preferred and non-preferred responses, as well as the difference between the responses to high-and low-valued objects (mean ± standard error).

After identifying individual value-coding neurons, we averaged the STN neuronal activities for each neuron’s preferred and non-preferred values to determine whether value representations were consistently expressed across the neuronal population (Fig. 3C, magenta and black lines, respectively). This approach enabled us to compare value selectivity across neurons with different value preferences. For flexible value, the non-preferred value response showed a sustained suppression, while the preferred value response exhibited a sustained excitation. This clear divergence between preferred and non-preferred value indicates that the flexible value is robustly represented across STN neurons. Similarly, stable value coding was also evident at the population level, although with distinct temporal dynamics. Both preferred and non-preferred value responses exhibited prominent phasic visual responses immediately after object presentation, followed by stronger activity for the preferred value than for the non-preferred value. Thus, despite their different response profiles, STN neurons discriminated both flexible and stable values. Collectively, these population-averaged responses demonstrate that value selectivity is consistently represented across STN neurons, while flexible and stable values exhibit distinct response dynamics.

### Stable value-coding neurons preferentially encode low-valued objects

Previous primate research demonstrated that the neurons in the GPe showed decreased activity to low-valued objects during stable value processing, suggesting that the indirect pathway preferentially represents stable value associated with habitual behavior^27^. Because the GPe and STN are reciprocally connected^48^, we asked whether the STN similarly exhibits a biased representation of stable value. To address this question, we first classified stable value-coding neurons according to their preference as positive coding neurons, which responded more strongly to high-valued objects, or negative coding neurons, which responded more strongly to low-valued objects. Among the 50 neurons encoding stable value (24 of stable value-coding neurons and 26 of dual value-coding neurons), 68% (n=34) were negative coding neurons and 32% (n=16) were positive coding neurons (Fig. 3A, lower right). Consistent with this population bias, standardized regression coefficients also showed that stable value-coding neurons were distributed preferentially toward negative coding (Fig. 3B). Furthermore, this bias was reflected in the population responses, with neuronal activity being stronger for low-valued than high-valued objects (Fig. 3D, right). Together, these results demonstrate that the STN represents stable value through negative coding, consistent with the biased representation previously reported in the GPe.

In contrast, flexible value coding showed a different pattern. Out of 73 neurons encoding flexible value (47 flexible value-coding neurons and 26 dual value-coding neurons), positive-coding (52%, n = 38) and negative-coding neurons (48%, n = 35) were almost equally represented (Fig. 3A, upper right). Likewise, standardized regression coefficients were distributed symmetrically along the flexible value axis (Fig. 3B), and averaged neuronal responses showed little difference between high-and low-valued objects (Fig. 3D, left). These findings indicate that, unlike stable value, flexible value is represented in a balanced manner without an apparent coding bias.

### Functional segregation of flexible and stable value coding along the rostral-caudal axis of the STN

We next asked whether distinct value-coding neurons are topographically organized within the STN. We therefore examined the anatomical distribution of flexible value-coding neurons (yellow), stable value-coding neurons (magenta), dual value-coding neurons (purple), and visual neurons (white) along the two-dimensional axes: rostral-caudal axis and dorsal-ventral axis (Fig. 4A). Interestingly, the STN neurons showed the most prominent segregation across the rostral-caudal axis. Flexible value-coding neurons were primarily distributed in the rostral part, while stable value-coding neurons were mainly located in the caudal part. We observed a gradual decrease in the proportion of flexible value-coding neurons, accompanied by an increase in the proportion of stable value-coding neurons across the STN from rostral to caudal subregions (Fig. 4A, lower left). Dual value-coding neurons and visual neurons exhibited similar proportions along the rostral-caudal and dorsal-ventral axes (Fig. 4A, lower left and upper right). We did not observe systematic anatomical segregation along the medial-lateral and dorsal-ventral axes (Fig. 4A, upper right, Fig. S2). These findings suggest distinct value processing in goal-directed and habitual behaviors specifically along the rostral-caudal axis.

**Figure 4.**
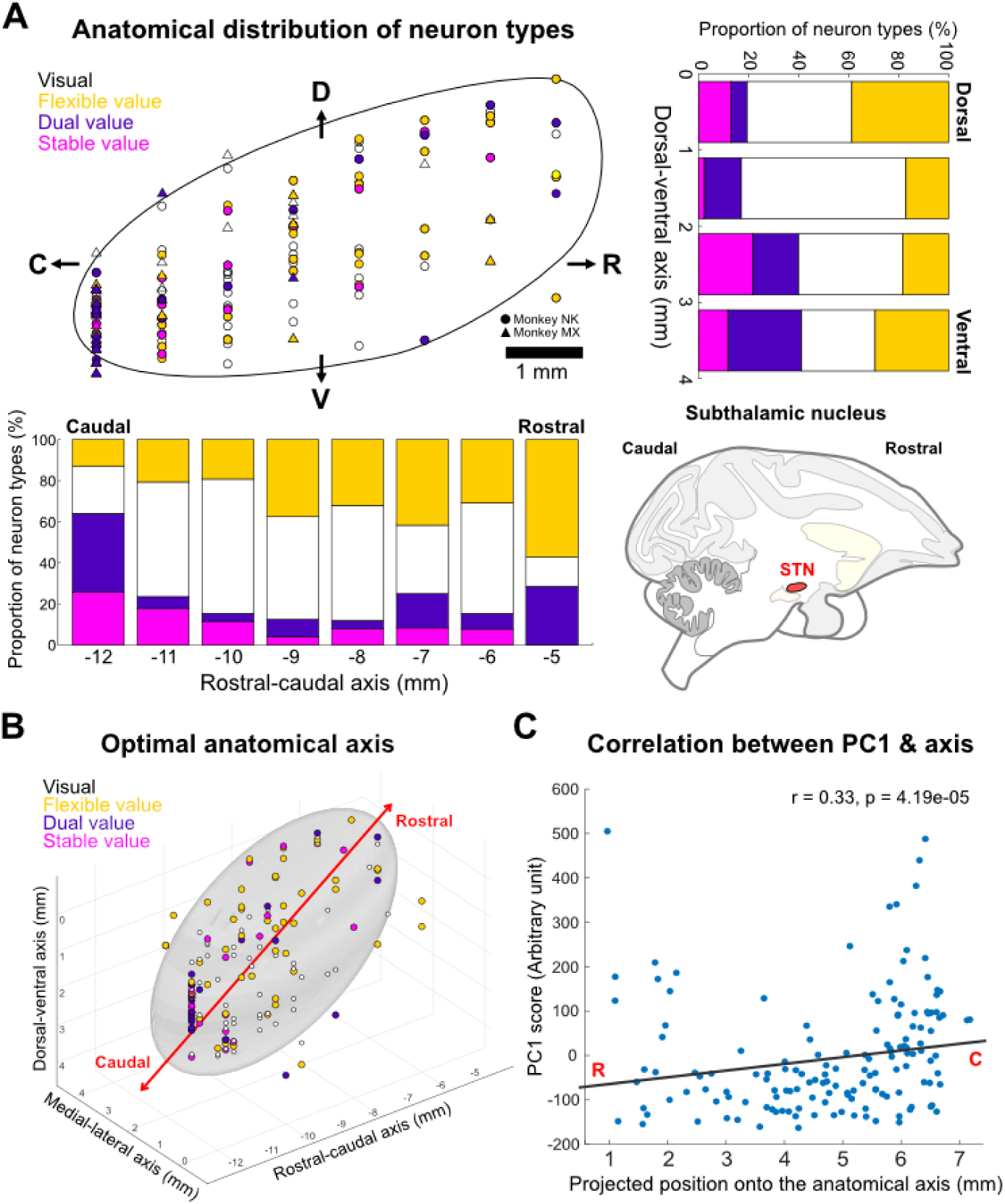
Distinct processing of flexible and stable values across the rostral-caudal axis in the STN. (A) Neuron type distribution along the anatomical axes. The upper left panel shows the structure of the STN and the anatomical distribution of neuron types. Neuron type distribution along the rostral-caudal axis (lower left panel) and dorsal-ventral axis (upper right panel). The white bar represents the portion of visual neurons, the yellow bar represents the portion of flexible value-coding neurons, the purple bar represents the portion of dual value-coding neurons, and the magenta bar represents the portion of stable value-coding neurons. The lower right panel illustrates the overall brain structure in a sagittal section, including the STN (red structure). (B) The best-fit axis (red arrow) across the STN that best corresponds to PC1. The recorded neurons are represented in the 3-dimensional anatomical structure of the STN. (C) Correlation between the neurons’ PC1 scores and their projected positions on the best-fit anatomical axis.

To further confirm the location of the recorded neuron within the STN, we made an electrolytic marking lesion and histologically reconstructed the location after recording an example dual-value coding neuron. Figure 5A shows the MR image used for targeting the cSTN. The marking lesion was identified within the cSTN using immunohistochemistry staining for vGLUT2 (vesicular glutamate transporter 2), a marker of glutamatergic neurons in the STN^49^ (Fig. 5B). Figure 5C illustrates the example dual-value coding neuron recorded from the cSTN at the site of the marking lesion. This neuron exhibited a strong inhibitory visual response to object presentation in both the flexible- and stable-value tasks. Following this initial inhibition, the neuron showed sustained inhibition in response to high-valued objects, whereas its activity rebounded and became excitatory in response to low-valued objects. Together, the anatomical reconstruction and neuronal response confirm that the recorded neuron was located within the cSTN and encoded both flexible and stable values.

**Figure 5.**
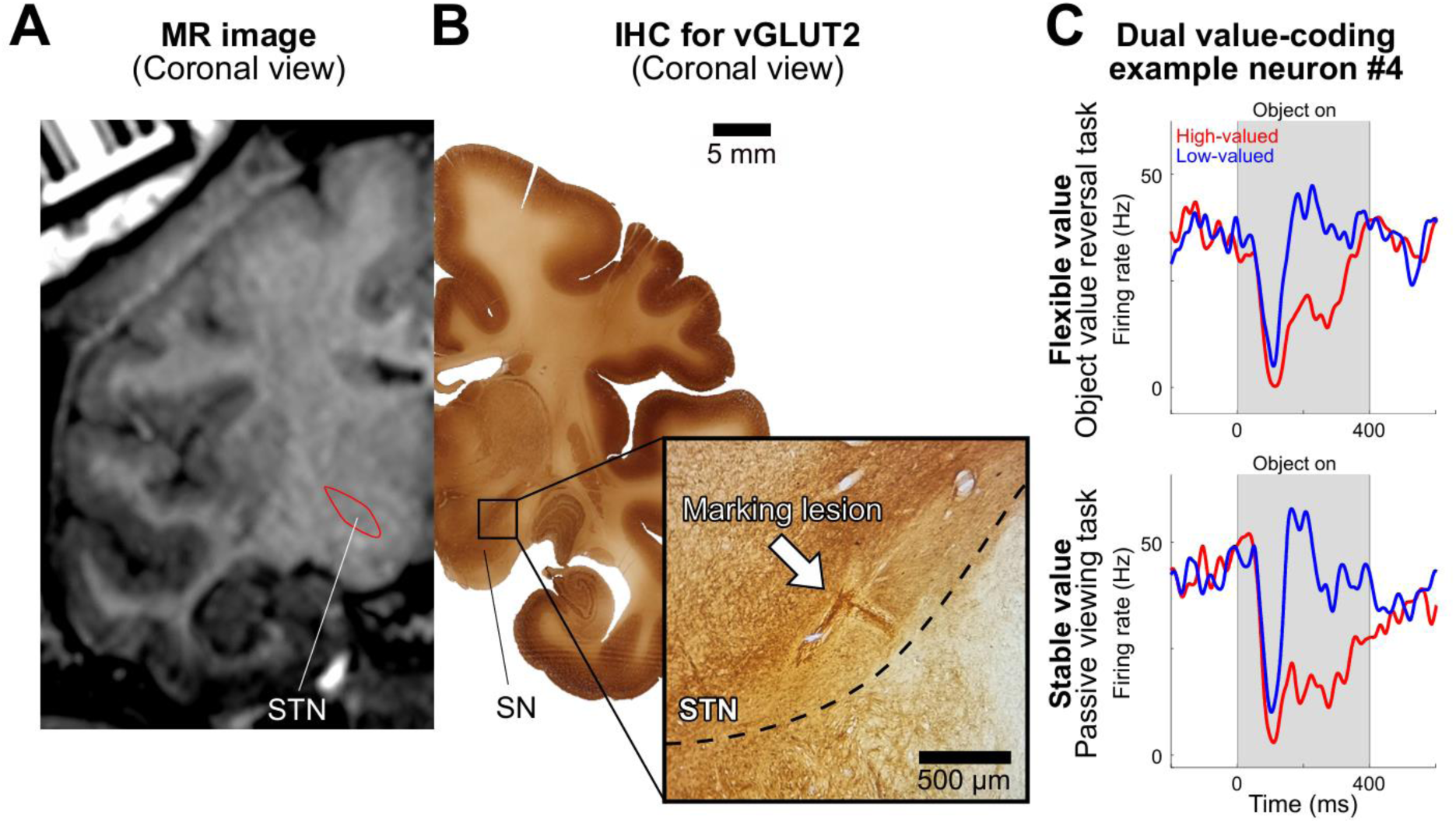
Coronal view of MRI and immunohistochemical images showing recording sites in the STN. (A) A magnetic resonance (MR) image in coronal view of the target recording sites. The STN was marked with a red outline. (B) Location of the STN neurons identified by immunohistochemistry (IHC) for vesicular glutamate transporter 2 (vGLUT2). Scale bar, 5 mm. The lower right inset shows the location of the STN neurons, indicated by a marking lesion (white arrow) in the immunohistochemistry image. The dashed black line indicates the ventral border of the STN. Scale bar, 500 μm. (C) Dual value-coding neuron recorded in the cSTN at the site of the marking lesion. The response of the example STN neuron to fractal objects is associated with flexible value in the object value reversal task (upper panel) and stable value in the passive viewing task (lower panel), and is aligned to the object onset. The red lines indicate the response to the high-valued objects, and the blue lines indicate the response to the low-valued objects.

To integrate anatomical and electrophysiological properties of the STN neurons, we conducted a principal component analysis (PCA) of the averaged z-scored neuronal activities per condition (flexible high-valued, flexible low-valued, stable high-valued, stable low-valued) and per time bin (every 100 ms from 200 ms before object onset to 600 ms after object onset). We first computed the scores of the 1st to 32nd principal components (PC1 to PC32) and projected neurons’ PC1 score onto all possible anatomical axes. Interestingly, PC1 described 80% of the variance in population neuronal activity.

Based on this analysis, we found the optimal axis by correlating the PC1 score with neurons’ projected position on an arbitrary anatomical axis^50^. We constructed a three-dimensional ellipsoid model of the STN and coordinates of recorded neurons (Fig. 4B). Among the tested axes, the axis shown in Figure 4B exhibited the strongest correlation between the PC1 score and neurons’ projected position on this axis. Therefore, we defined this axis as the optimal axis. Figure 4C depicts the significant correlation between PC1 scores and projected anatomical positions (r = 0.33, p = 4.19e-05, Spearman correlation), which implies that the neuronal response pattern of the STN is largely determined by the anatomical organization. The best-fit axis suggests that the STN shows distinct neuronal properties along the rostral-caudal axis, as also indicated by the anatomical distribution. Overall, these findings reveal a gradual shift in value representation, from predominantly flexible value coding rostrally to increasingly stable value coding caudally.

### Distinct behavioral effects of rSTN and cSTN inactivation on goal-directed and habitual decision-making

We revealed a gradual functional organization in STN neuronal activity along the rostral-caudal axis, with flexible and stable value signals represented in distinct subregions. A critical unresolved question, however, is whether this anatomical organization merely reflects neuronal coding or causally supports parallel control of distinct value-based behaviors.

To determine whether their distinct value representations causally contribute to goal-directed and habitual behaviors, we separately inactivated the rSTN and cSTN. We injected muscimol (gamma-aminobutyric acid (GABA) agonist) into the rSTN or cSTN to achieve region-specific inactivation (Fig. 6C). Before drug injection, we first identified the target subregion by recording neuronal activity using an injectrode, which enables both extracellular recording and local drug delivery. Once the target neuron was identified, muscimol was injected, and the monkey performed the goal-directed value reversal choice task to evaluate flexible value-guided behavior and the habitual free looking task to assess stable value-guided behavior. We obtained pre-and post-muscimol injection behavioral data, with the pre-injection data serving as the control condition.

**Figure 6.**
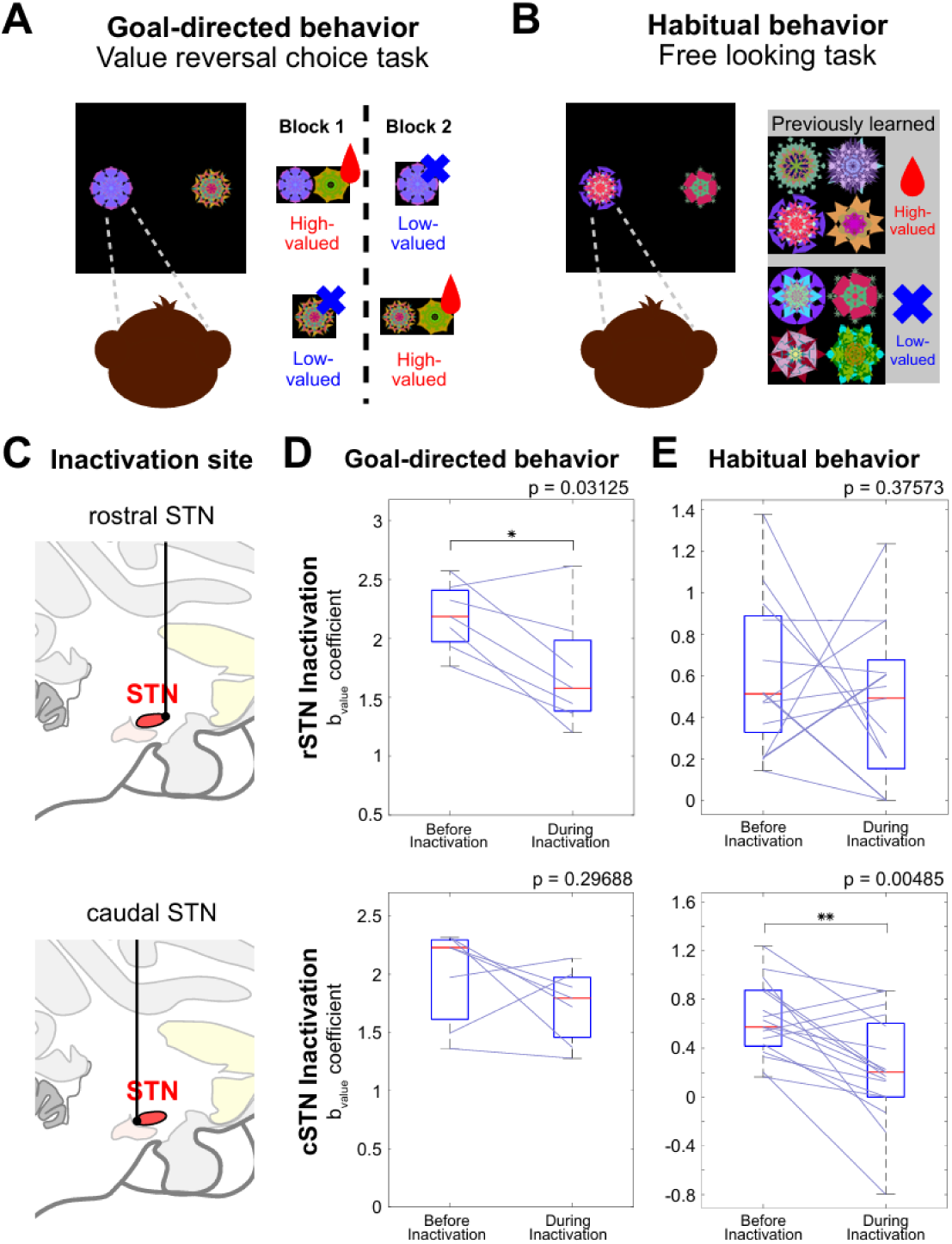
Differential impairments of value-based goal-directed and habitual behaviors following rSTN and cSTN inactivation with muscimol. (A) Value reversal choice task procedure for testing the inactivation effect on goal-directed behavior. One high-valued object and one low-valued object were presented simultaneously, one on the left side and the other on the right side. The object-reward contingency was changed every 24 trials. The monkey was forced to make a choice between two objects. After the monkey chose the high-valued object, the monkey received the reward. (B) Free looking task procedure for testing the inactivation effect on habitual behavior (same as in Figure 1C). (C) Example injection sites of muscimol in the STN (red structure): rostral STN (upper panel) and caudal STN (lower panel). (D and E) Inactivation effect on value-based goal-directed behavior (D) and habitual behavior (E) following rSTN inactivation (upper panels) and cSTN inactivation (lower panels). Each line indicates the pair of value coefficients obtained from logistic regression before and during inactivation. The red line in the blue box represents the mean, and the blue box corresponds to the distribution from the 25th to 75th percentiles. *p < 0.05, **p < 0.005.

In the value reversal choice task to assess goal-directed behavior, a high-valued object and a low-valued object were presented on opposite sides of the screen, with their left-right positions pseudorandomized across trials. The monkey had to make a saccadic choice between two objects. The reward was delivered only if the monkey chose the high-valued object (Fig. 6A). These objects changed their object-reward association every 24 trials, requiring the monkey to update flexibly changing object values and engage in goal-directed behavior. To assess habitual behaviors, the monkey performed the same free looking task from Fig. 1C (Fig. 6B). In this task, previously learned high-and low-valued objects were presented without any reward outcome, with spontaneous orienting toward high-valued objects reflecting stable value-driven habitual behaviors.

First, we compared correct rate and habitual-choice rate before and during STN inactivation while the monkey performed value reversal choice and free looking tasks. Correct rate reflects the proportion of trials in which the monkey selected the flexible high-valued object during goal-directed behavior, whereas habitual-choice rate reflects the degree to which the monkey preferentially chose the stable high-valued object over the stable low-valued object during habitual behavior. Following rSTN inactivation, the correct rate significantly decreased (Fig. S3A, top) (p = 0.03125, Wilcoxon signed-rank test), and habitual-choice rate showed a non-significant trend toward a decrease (Fig. S3B, top) (p = 0.09228, Wilcoxon signed-rank test). Conversely, cSTN inactivation significantly reduced the habitual-choice rate (Fig. S3B, bottom) (p = 8.38e-4, Wilcoxon signed-rank test), while the correct rate remained unchanged (Fig. S3A, bottom) (p = 0.23438, Wilcoxon signed-rank test). These findings are consistent with our electrophysiological results showing that the rSTN preferentially processes flexible value, whereas the cSTN preferentially processes stable value.

However, because both the value reversal choice task and the free looking task involved binary choices, behavioral performance could be influenced not only by object value but also by other factors, such as spatial preference and stimulus salience. In particular, previous studies have reported that STN inactivation can bias contralateral eye movements^51–53^. Thus, a directional bias toward the contralateral side could reduce correct rate or habitual-choice rate, making it difficult to distinguish a disruption of value-guided behavior from a change in spatial preference. In addition, stimulus salience affects visual choice behavior^54–56^. Therefore, to quantify the contribution of value to choice behavior while accounting for other factors that could influence the monkey’s choices, we fitted a logistic regression model to the behavioral data. The model included coefficients for object value (b_value_), spatial direction (b_direction_), and stimulus salience (b_saliency_). The value coefficient quantified the extent to which the relative value of the two objects influenced the probability of choosing one object over the other, the direction coefficient captured the influence of spatial preference on choice, and the salience coefficient quantified the influence of the relative salience of the two objects on choice. Thus, changes in the value coefficient after STN inactivation provided a measure of how strongly choice behavior remained guided by object value, independent of changes in direction-or salience-related biases. We then compared these coefficients between the pre-and post-injection conditions to determine whether rSTN or cSTN inactivation selectively altered value-guided goal-directed or habitual behavior.

Under rSTN inactivation, the b_value_ coefficient decreased while performing the goal-directed value reversal choice task (Fig. 6D, top) (p = 0.03125, Wilcoxon signed-rank test). These results suggest that rSTN inactivation weakened value-guided choice behavior in the reversal task. In contrast, the monkey showed a non-significant change in the b_value_ coefficient after rSTN inactivation (Fig. 6E, top) (p = 0.37573, Wilcoxon signed-rank test). This indicates no effect on value-based habitual behavior, implying intact habitual behavior after the rSTN inactivation. Taken together, these results indicate that the rSTN contributes only to goal-directed behavior.

Following cSTN inactivation, the monkey chose the high-valued object equally in the value reversal choice task (p = 0.29688, Wilcoxon signed-rank test). The b_value_ coefficient in the value reversal choice task remained unchanged (Fig. 6D, bottom), indicating that the cSTN does not play a role in goal-directed behavior. Conversely, the b_value_ coefficient in the free looking task decreased, indicating that the monkey made fewer automatic saccades toward high-valued objects (Fig. 6E, bottom) (p = 0.00485, Wilcoxon signed-rank test). This implies that the cSTN plays a role in habitual behavior based on stable value. In brief, the rSTN inactivation led to a decreased b_value_ coefficient during only goal-directed behavior, and cSTN inactivation caused fewer habitual and automatic saccades toward high-valued objects. These results indicate exclusive functions of goal-directed and habitual behaviors in the rSTN and cSTN, respectively (Fig. 6D-E). The inactivation results are consistent with the anatomical distribution of value-processing neurons in the STN along the rostral-caudal axis (Fig. 4A). This double dissociation implies the parallel circuit of value-based behavior control across the rostral-caudal axis in the STN. Moreover, we demonstrated that the STN not only processes flexible and stable value, but also causally contributes to value-based behavior.

In addition, we measured changes in the b_direction_ and b_saliency_ coefficients before and during inactivation to quantify the spatial direction and stimulus saliency to choice behavior. Following both rSTN and cSTN inactivation, the b_saliency_ coefficient during either goal-directed or habitual behaviors did not show significant changes (Fig. S4A). These findings suggest that the STN does not contribute to saliency-related eye movements. In contrast, regardless of which STN subregion was inactivated, only the b_direction_ coefficient during habitual behavior was significantly reduced (Fig. S4B, right panels), while the b_direction_ coefficient during goal-directed behavior remained unchanged (Fig. S4B, left panels). This indicates an increased preference for the contralateral direction, consistent with previous reports of contralateral choice biases following inactivation of the STN^51–53^. Notably, this directional bias was observed during habitual behavior. Because the habitual task involved spontaneous eye movements in the absence of an explicit task goal or instructed target, these results suggest that directional biases induced by STN inactivation are more readily expressed during unconstrained habitual eye movements. Collectively, we found that STN inactivation preferentially alters directional and value-related influences on eye movements rather than saliency-based influences.

## Discussion

To account for these findings, we propose a circuit model for the value processing mechanism in the STN. The schematics illustrate two distinct circuits guiding goal-directed and habitual behaviors in parallel. The caudate nucleus (CD), which receives dopaminergic input from the SNc, sends value information to the GPe. The GPe conveys value information to the STN via the loop circuit between the STN and GPe, and STN neurons transmit excitatory input to the SNr. The SNr inhibits neuronal activity in the superior colliculus (SC), thereby contributing to value-based eye movements toward high-valued objects.

Specifically, the caudal stream from the caudate tail (CDt) to cSTN regulates habitual behavior, and the rostral stream from the caudate head (CDh) to rSTN controls goal-directed behavior (Fig. 7). As previously described, we found that stable value-coding neurons preferentially responded to low-valued objects, whereas flexible value-coding neurons were evenly distributed between positive and negative coding types. The majority of the stable value-coding neurons responded more strongly to low-valued objects. This excitation leads to increased responses to low-valued objects in the caudal-dorsal-lateral SNr (cdlSNr). Consequently, the cdlSNr sends inhibitory GABAergic value information to the SC, which is the final output structure for eye movement control. Therefore, the neurons in the SC fire more frequently in response to stable high-valued objects, leading to saccadic eye movements toward them.

**Figure 7.**
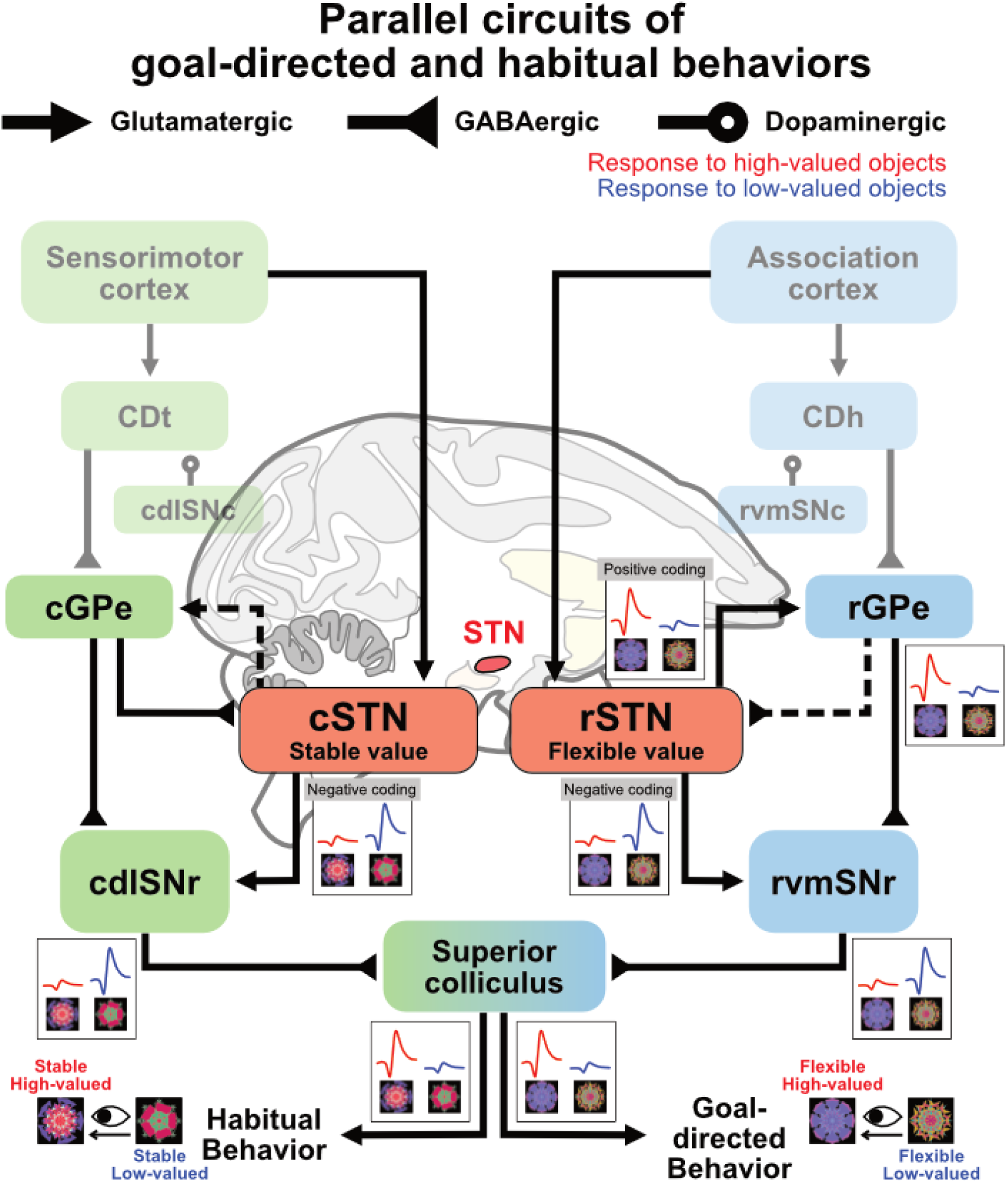
Parallel circuits of goal-directed and habitual behaviors associated with the rSTN and cSTN. Parallel circuits of goal-directed and habitual behaviors. The red lines indicate the conceptual responses to the high-valued objects, and the blue lines indicate the conceptual responses to the low-valued objects. CDt, caudate tail. CDh, caudate head. cdlSNc, caudal-dorsal-lateral substantia nigra pars compacta. rvmSNc, rostral-ventral-medial substantia nigra pars compacta. cGPe, caudal globus pallidus externa. rGPe, rostral globus pallidus externa. cSTN, caudal STN. rSTN, rostral STN. cdlSNr, caudal-dorsal-lateral substantia nigra pars reticulata. rvmSNr, rostral-ventral-medial substantia nigra pars reticulata.

On the other hand, for goal-directed behaviors guided by flexible value, flexible value-coding neurons were evenly distributed between positive and negative coding types. Half of the flexible value-coding neurons in the STN responded more strongly to high-valued objects and the other half responded more strongly to low-valued objects. Positive and negative coding type neurons convey flexible value information to the rostral GPe (rGPe) and the rostral-ventral-medial SNr (rvmSNr), respectively. Since the rGPe sends inhibitory GABAergic inputs and the rSTN transmits excitatory glutamatergic inputs to the rvmSNr, the rvmSNr is also excited by low-valued objects. In turn, the SC fires more frequently to the flexible high-valued object, guiding goal-directed eye movements toward flexible high-valued objects (Fig. 7).

Consistent with this framework, inactivation of the rSTN and cSTN resulted in impairments in goal-directed and habitual decision-making, respectively. The b_value_ coefficient in the reversal choice and free looking tasks decreased after rSTN and cSTN inactivation, respectively (Fig. 6D-E). These findings indicate that value signals encoded in distinct STN subregions contribute causally to different forms of value-based behavior.

Together, our results suggest that the STN encodes both flexible and stable values in a parallel manner, with these signals organized along the rostral-caudal axis. The rSTN is more closely associated with flexible value processing, whereas the cSTN is linked to stable value processing. This organization is consistent with functional principles observed in other basal ganglia structures and supports a circuit-level framework in which goal-directed and habitual behaviors are mediated by partially segregated pathways.

From a clinical perspective, these findings may provide insight into cognitive dysfunctions in Parkinson’s disease. As the STN contributes to value-guided decision-making based on flexible and stable values, alterations in subthalamic activity associated with PD or DBS might disrupt the balance of value signals across downstream circuits and affect value-based decision-making. In this context, the rostral-caudal organization of the STN is relevant for understanding variability in behavioral outcomes following STN-DBS. Thus, the anatomical axis delineating value processing across the STN could guide more efficient, region-specific DBS protocols and establish the general relationship between the STN-DBS regions and their clinical effects.

In conclusion, consistent with other basal ganglia structures, the STN exhibits a similar functional organization: the rostral stream supporting cognitive flexibility and the caudal stream supporting habitual stability. These findings provide a circuit-level framework linking STN activity to value-based behavior and offer a basis for future studies investigating the neural mechanisms of cognitive dysfunction in Parkinson’s disease.

## Methods

### Animals

One male and one female rhesus monkey (*Macaca mulatta*, 8.0 and 5.0 kg, referred to as MX and NK) were used. Animal care and experimental procedures were performed as approved by the Seoul National University Institutional Animal Care and Use Committee. Under general anesthesia and surgical conditions, a plastic head holder and a recording chamber were implanted into the monkeys’ skulls. The position of each chamber was tilted laterally by 25° such that it corresponded to the striatum. We started the training and recording sessions after the monkeys fully recovered from surgery.

### Behavioral tasks

The behavioral procedures were controlled by a custom-made Visual C^++^-based experimental data acquisition system (Blip; available at http://www.cocila.net/blip/). The monkey sat in a primate chair, facing a frontoparallel monitor screen in a sound-attenuated and electrically shielded room. Visual fractal stimuli created using Fractal Geometry were approximately 8° x 8° in size. The mean luminance was equalized across the images using the Spectrum, Histogram, and Intensity Normalization and Equalization (SHINE) toolbox written in MATLAB. An infrared camera (Oculomatic Pro) was used to record the eye position with a sampling rate of 1 kHz.

### Object value reversal task

To examine behavioral performance and neuronal encoding of flexibly changing object value, we performed a task in which the object-value contingency was reversed in each block of 20–25 trials^32,42^. For each monkey, fixed sets of two fractal objects (36 sets for monkey MX and 15 sets for monkey NK) were used as the saccade targets. In each trial, one of them was presented at the right or left position pseudo-randomly (15° from the center). The fixation dot was presented simultaneously with the fractal object and disappeared after a 400-ms overlap period for recording the neural responses to the objects. Following the disappearance of the fixation dot, the monkeys were required to select the presented object by making a saccade to proceed to the next trial. In a block of 20–25 trials, one of the objects was associated with a reward (high-valued object) and the other with no reward (low-valued object). In the next block, the object-reward contingency was reversed. At least four blocks were included in one experiment. The object value reversal task for the monkey NK included choice trials wherein the two objects were simultaneously presented at the right and left positions, and the monkey had to choose between the two objects. If the monkey correctly selected the high-valued object, a liquid reward was provided. The choice trial appeared every fifth trial.

### Value reversal choice task

To examine the causal contribution of the STN to value processing, we performed a task composed solely of choice trials before and after muscimol injection. One high-valued and one low-valued object were presented at the right and left positions, and if the monkey correctly selected the high-valued object, a liquid reward was delivered. To increase task difficulty and prevent context-dependent choices, among the three objects, either two were high-valued and one was low-valued, or one was high-valued and two were low-valued. The object-value contingency was reversed in each block of 24 trials, and at least four blocks were included in one experiment.

### Stable value learning task

To generate stable long-term value memory, monkeys viewed visual objects repeatedly in association with consistent reward outcomes and thereby learned their stable values^32,42,57^. In each of these sessions and the following tasks, a set of 8 computer-generated fractals was employed as visual objects. The stable value learning task included 64 forced trials and 32 choice trials. In forced trials, while the monkey was fixating on a central white dot, one of the objects was presented at the ipsilateral or contralateral position (15° from the center). The central fixation spot turned off 400 ms later, and the monkey was required to make a saccade to the object. Half of the objects were associated with a liquid reward (high-valued objects), whereas the other half were associated with no reward (low-valued objects). A tone was presented with either outcome. In choice trials, one high-valued and one low-valued object were simultaneously presented at the right and left positions, and the monkey had to choose between the two objects. If the monkey correctly selected the high-valued object, a liquid reward was provided. The choice trial appeared every third trial. Each set of objects was learned in one learning session per day. The same sets of objects were repeatedly learned with the same object-value associations across days (>4 days), while new sets of fractals were introduced for learning across days. The monkeys MX and NK learned 51 (408 objects) and 73 (584 objects) sets of objects, respectively. Different sets of fractal objects were employed in procedures for testing cognitive flexibility and habitual stability.

### Free looking task

To confirm whether the monkeys displayed a visual habit based on stable object value, we tested free gaze during which no instruction was provided, but the learned fractal objects were presented, as previously used in monkeys and humans^42,57^. In a free looking task with two objects, one high-valued and one low-valued object were selected pseudo-randomly from a set of eight learned objects and presented simultaneously at the right and left positions (15° from the center) after the monkey had fixated on a central white dot for 300 ms. The monkey was free to look at them for 2 s without any reward outcome. In addition to the free looking trials, a reward-associated white dot was presented during half of the trials to maintain the monkey’s motivation. In the free looking task with two objects, this dot was presented at one of three positions (i.e., the top, middle, and bottom). The reward was delivered when the monkey held its gaze on the white dot for 600 ms.

### Passive viewing task

This task was used to examine how neurons responded to the previously learned objects, as previously described^32,42,57^. While the monkey was fixating on a central white dot, two to four objects, selected pseudo-randomly from a set of eight learned objects, were sequentially presented for 400 ms each at 15° from the center. Each learned object was presented at least 10 times in one session. To maintain the monkeys’ motivation, the reward was delivered 300 ms after the last object disappeared. Thus, the reward was not directly associated with any object. The value-coding activity was recorded after long-term learning (>4 days) with a sufficient retention period (>1 day after the last learning session).

### Single-unit recording

As a monkey performed a task, the activity of single neurons in the subthalamic nucleus was recorded using a general method. The recording sites were determined by means of a 1-mm spacing grid system with the aid of MR images (3T, Siemens) obtained along the direction of the chamber. The single-unit recording was conducted using a glass-coated electrode (Alpha-Omega). The electrode was inserted into the brain through a stainless-steel guide tube and advanced by an oil-driven micromanipulator (MO-974A, Narishige). Neuronal signals from the electrode were amplified, filtered (250 Hz to 10 kHz), and digitized (30-kHz sampling rate and 16-bit A/D resolution) using a Scout system (Ripple Neuro, UT). Neuronal spikes were isolated online using custom voltage-time window-discrimination software (BLIP, Laboratory of Sensorimotor Research, National Eye Institute, National Institutes of Health [LSR/NEI/NIH], available at www.cocila.net/blip), with the corresponding timings detected at 1 kHz. The waveforms of individual spikes were collected at 50 kHz.

### Analysis of object response

To examine the neuronal object responses, we counted the number of spikes within the test and control windows for each fractal object in the passive viewing task and object value reversal task. The control window was defined as 0–200 ms before the object onset, and the test window was defined as 0–400 ms after the object onset. To test whether the neurons showed object responses, we compared the numbers of spikes between the control and test windows in individual trials for each object. The Wilcoxon rank-sum test was employed to test for statistical significance.

### Analysis of value-coding activity

We performed linear regression analysis to determine whether the discharge rates of individual neurons recorded during the test window, as previously described, were significantly related to the value of an object. The discharge rates during the test window (*Y*) were fitted by the following linear regression model (Equation 1).

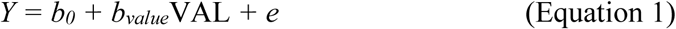

where VAL was the value of an object (1 and -1 for high value and low value); *b_value_* was the regression coefficient; and *b_0_* and *e* were the intercept and residual, respectively. If *b_value_* was significantly different from zero (P < 0.05, t-test) and positive, we regarded the neuron as a positive value-coding neuron. If the *b_value_* was significantly different from zero (P < 0.05, t-test) and negative, we regarded the neuron as a negative value-coding neuron. Both positive and negative value-coding neurons were classified as value-coding neurons.

### Histology, marking lesion, and immunohistochemistry

Monkey MX was deeply anesthetized with an overdose of sodium pentobarbital and perfused transcardially with saline followed by 4% paraformaldehyde. The head was fixed to the stereotaxic frame, and the brain was cut into blocks in the coronal plane, including the midbrain region. The block was post-fixed overnight at 4℃, and then cryoprotected for 5 days in increasing gradients of glycerol solution (5, 10, 20% glycerol in PBS) before being frozen. The frozen block was cut every 50-μm using a microtome.

To histologically confirm and reconstruct the locations of putative STN neurons, after recording the value-coding neurons in the STN, we passed a 13 μA-negative current for 30 s to make a marking lesion. To test whether this marking lesion and the recorded neuron were located in the STN, we labeled 600 μm-interval STN slices with a vGLUT2 (vesicular glutamate transporter 2) antibody. After 1 h of permeabilization with 0.05% Triton X-100 in PBS (phosphate-buffered saline), the slices were blocked for 1 h in a solution containing 3% normal goat serum, 2% bovine serum albumin, and 0.05% Triton X-100 in PBS. The blocked slices were incubated with a mouse anti-vGLUT2 (1:1000; Sigma-Aldrich, MAB5504) antibody overnight at room temperature. The next day, the slices were washed with PBS, and then incubated for 1 h at room temperature with anti-mouse biotinylated secondary antibody and prepared peroxidase conjugate (Vectorlabs, PK-6102). After the staining procedure, the slices were air-dried overnight at room temperature.

### Inactivation of the subthalamic nucleus

To inactivate each region of the STN, we injected muscimol (GABA_A_ agonist) into the rostral STN or caudal STN. The injection was performed on the right side of the STN of the monkey. To accurately locate the injection site, we recorded single-unit activity before the injection. For this purpose, we used a custom-made injectrode consisting of an epoxy-coated tungsten microelectrode for neuron recording and a silica tube for muscimol injection. Before the injection, while the injectrode was positioned at the injection site, the monkey performed the value reversal choice task and free looking task, and the data were used as a pre-injection control. We injected 0.6-1.2 μl of 5.12 mM muscimol (Sigma-Aldrich) at a speed of 0.2 μl/min. Starting 5 min after the injection, the monkey was required to resume the flexible and stable value tasks. The tests were repeated several times until 2-3 h after the injection. We performed the inactivation experiments after collecting most of the behavioral and neuronal data.

### Analysis of inactivation study

To examine value-based behaviors before and after the inactivation of the STN, we performed the logistic regression analysis. The choice of contralateral object (Y) was fitted by the following logistic regression model (Equation 2).

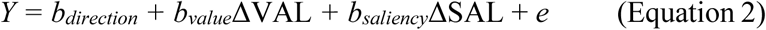

where ΔSAL was the difference between the saliency of the contralateral and ipsilateral objects (ΔSAL = 1 if the ipsilateral object was high-saliency, and ΔSAL = −1 if the contralateral object was high-saliency); ΔVAL was the difference between the values of the contralateral and ipsilateral objects (ΔVAL = 1 if the ipsilateral object was high-valued, and ΔVAL = −1 if the contralateral object was high-valued); *b_saliency_* was the regression coefficient; *b_value_* was the regression coefficient; *b_direction_* was the intercept indicating the bias toward choosing the contralateral object; and *e* was the residual.

Then, we compared *b_saliency_, b_value_* and, *b_direction_* before and after the inactivation of the STN to determine whether the behavior was compromised after the inactivation of the STN. If the *b_value_* decreased significantly after the inactivation (P < 0.05, Wilcoxon signed-rank test), we considered that the inactivation impaired value-based behavior. If the *b_saliency_* increased or decreased after the inactivation (P < 0.05, Wilcoxon signed-rank test), we considered that the inactivation altered saliency-based behavior. If the *b_direction_* increased or decreased after the inactivation (P < 0.05, Wilcoxon signed-rank test), we considered that the inactivation caused ipsilateral or contralateral preference, respectively.

## Acknowledgments

This work has been supported by the Bio&Medical Technology Development (RS-2025-02263832), the Basic Science Research (RS-2024-00339355), the ASTRA (RS-2024-00436783) through the National Research Foundation of Korea (NRF) funded by the Korean government (MEST), the LAMP (RS-2023-00301976) by the Ministry of Education, and the IITP (RS-2026-25507282) funded by the Korea government (MSIT).

## Author contributions

H.F.K. designed and supervised the entire project. KW. L., M. L., H. J., and H.F.K. performed the behavior and single-unit recording experiments. KW. L., H. J., and H.F.K. analyzed the data and prepared the figures. KW. L. and H.F.K. wrote the first draft, and KW. L. and H.F.K. interpreted data and wrote the final manuscript.

## Competing interests

The authors declare that they have no competing interests.

## Data availability

All data needed to evaluate the conclusions in the paper are present in the paper and/or the Supplementary Materials.

The original/source data can be provided by Hyoung F. Kim pending scientific review and a completed material transfer agreement. Requests for the original/source data should be submitted to: Hyoung F. Kim.

---

**Figure S1.**
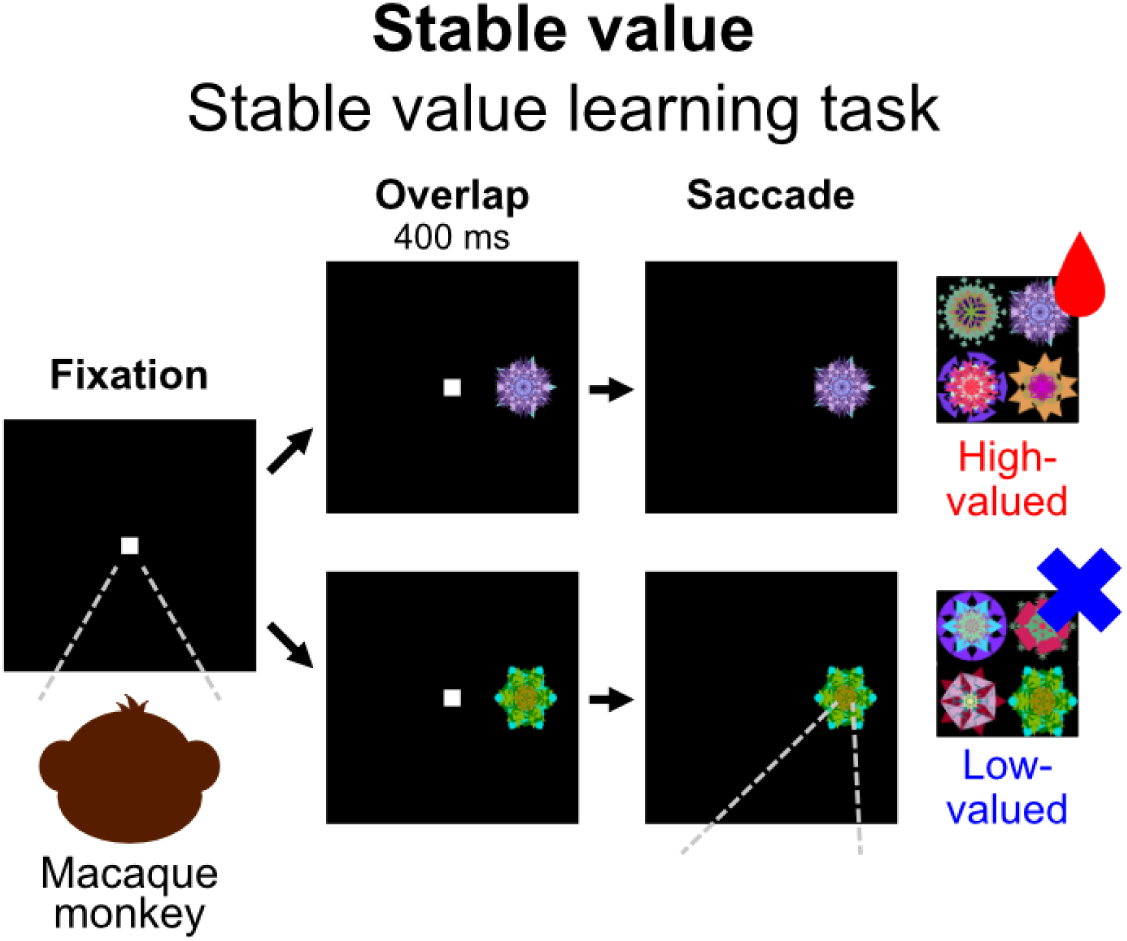
Stable value learning procedure. The scheme of the stable value learning task for learning stable value and testing habitual behavior. While the monkey was fixating on a central fixation dot, one of the eight objects appeared at the left or right position. After 400 ms, as the central fixation dot disappeared, the monkey was forced to make a saccade toward the object. 4 objects are associated with reward (high-valued), and the other 4 objects are not associated with reward (low-valued). The monkey received liquid reward only when the monkey made a saccade toward the high-valued object. This stable value learning procedure was repeated across >4 days.

**Figure S2.**
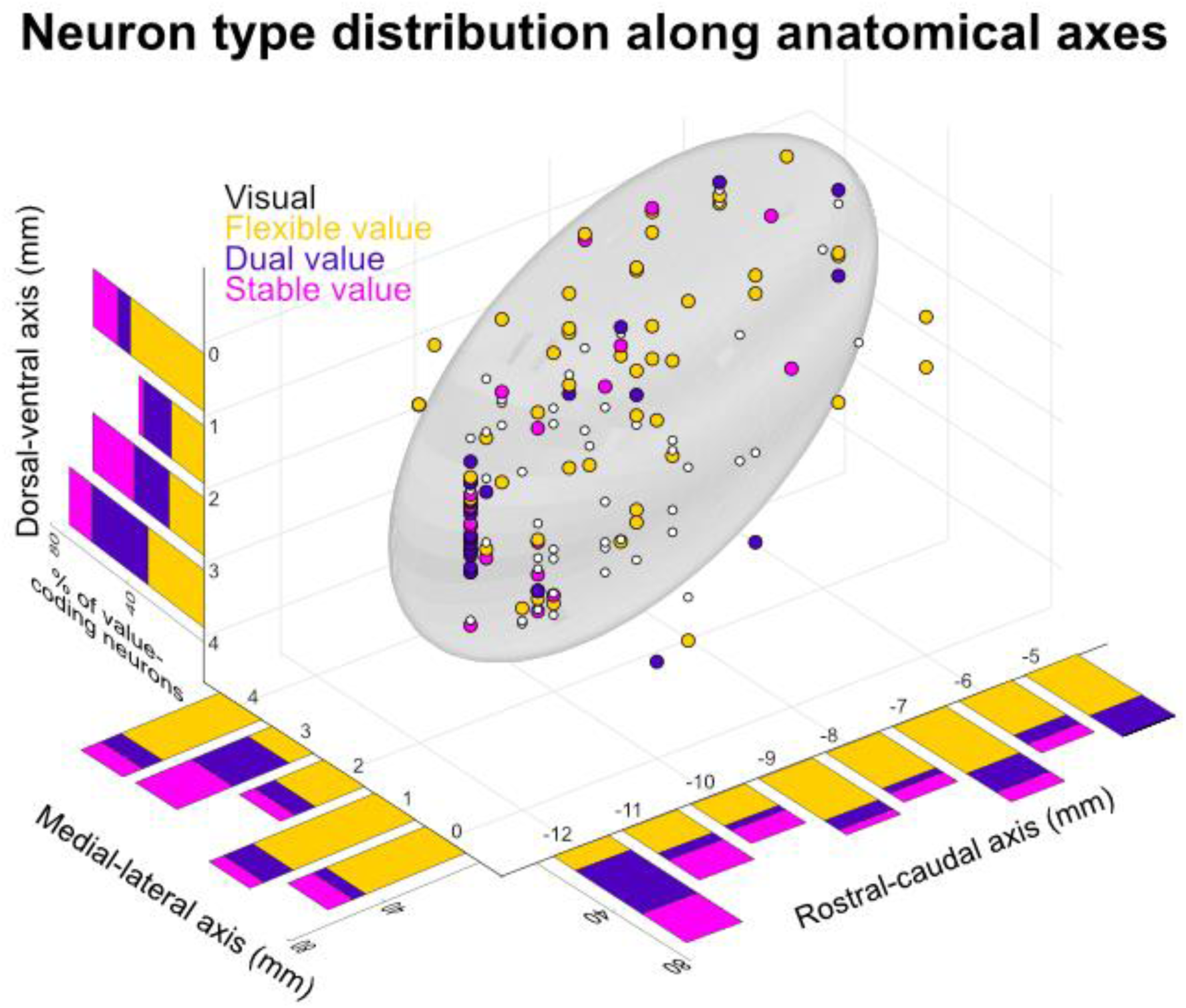
Neuron type distribution along anatomical axes. Neuron type distribution along the dorsal-ventral axis (left), medial-lateral axis (lower left), and rostral-caudal axis (lower right). The yellow bar represents the portion of flexible value-coding neurons, the purple bar represents the portion of dual value-coding neurons, and the magenta bar represents the portion of stable value-coding neurons. In the 3-dimensional structure of the STN (right ellipsoid), the white dots represent the visual neurons, the yellow dots represent the flexible value-coding neurons, the purple dots represent the dual value-coding neurons, and the magenta dots represent the stable value-coding neurons.

**Figure S3.**
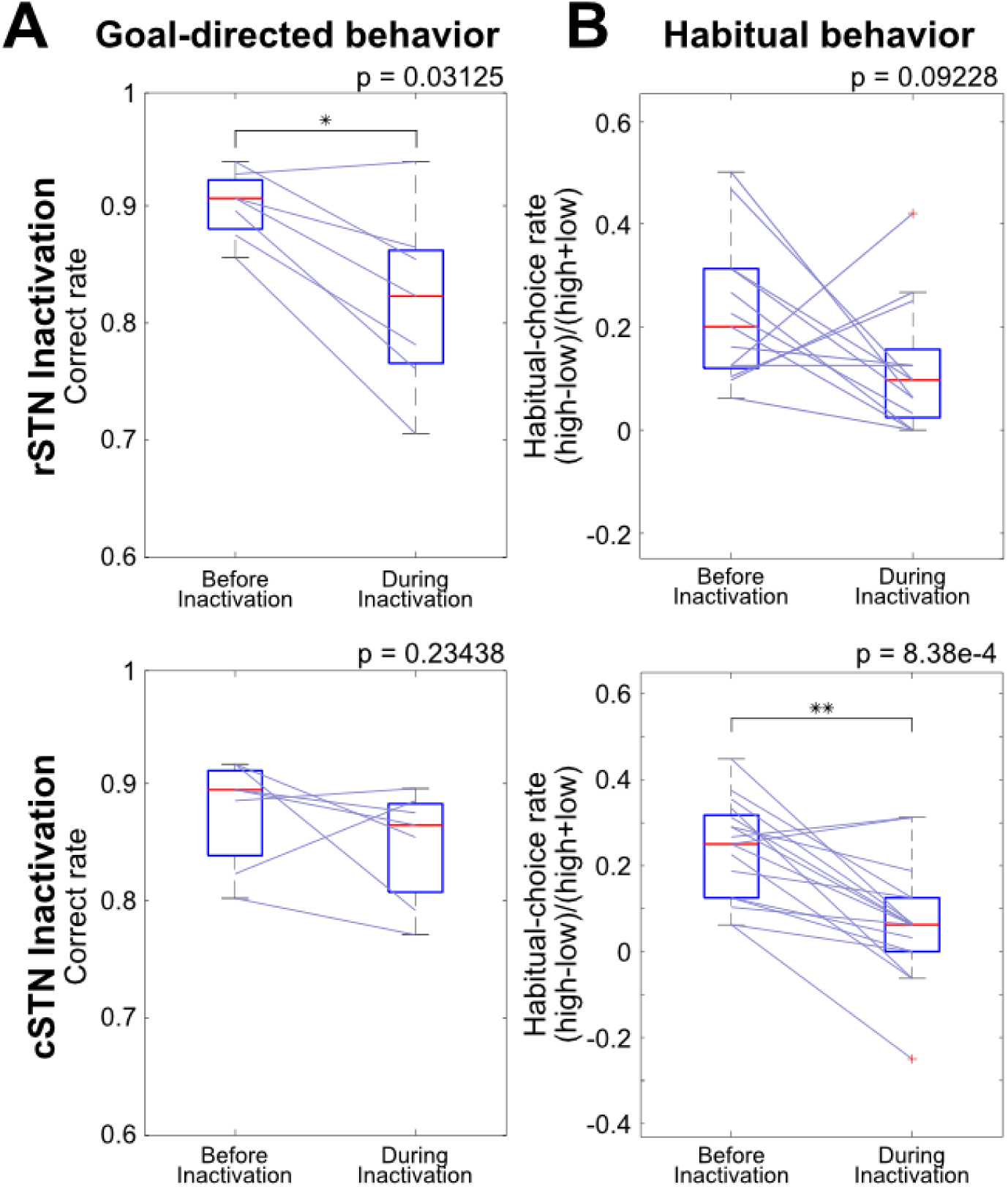
Impairments of goal-directed and habitual behaviors revealed by raw behavioral measures following rSTN and cSTN inactivation. (A and B) Inactivation effect on value-based goal-directed behavior (A) and habitual behavior (B) revealed by raw behavioral measures following rSTN inactivation (upper panels) and cSTN inactivation (lower panels). The correct rate (A) was defined as (high) / (high + low), and the habitual-choice rate (B) was defined as (high -low) / (high + low), with ‘high’ and ‘low’ referring to the number of saccades toward high-valued and low-valued objects, respectively. Each line indicates the pair of behavioral parameters before and during inactivation. The red line in the blue box represents the mean, and the blue box corresponds to the distribution from the 25th to 75th percentiles. *p < 0.05, **p < 0.005.

**Figure S4.**
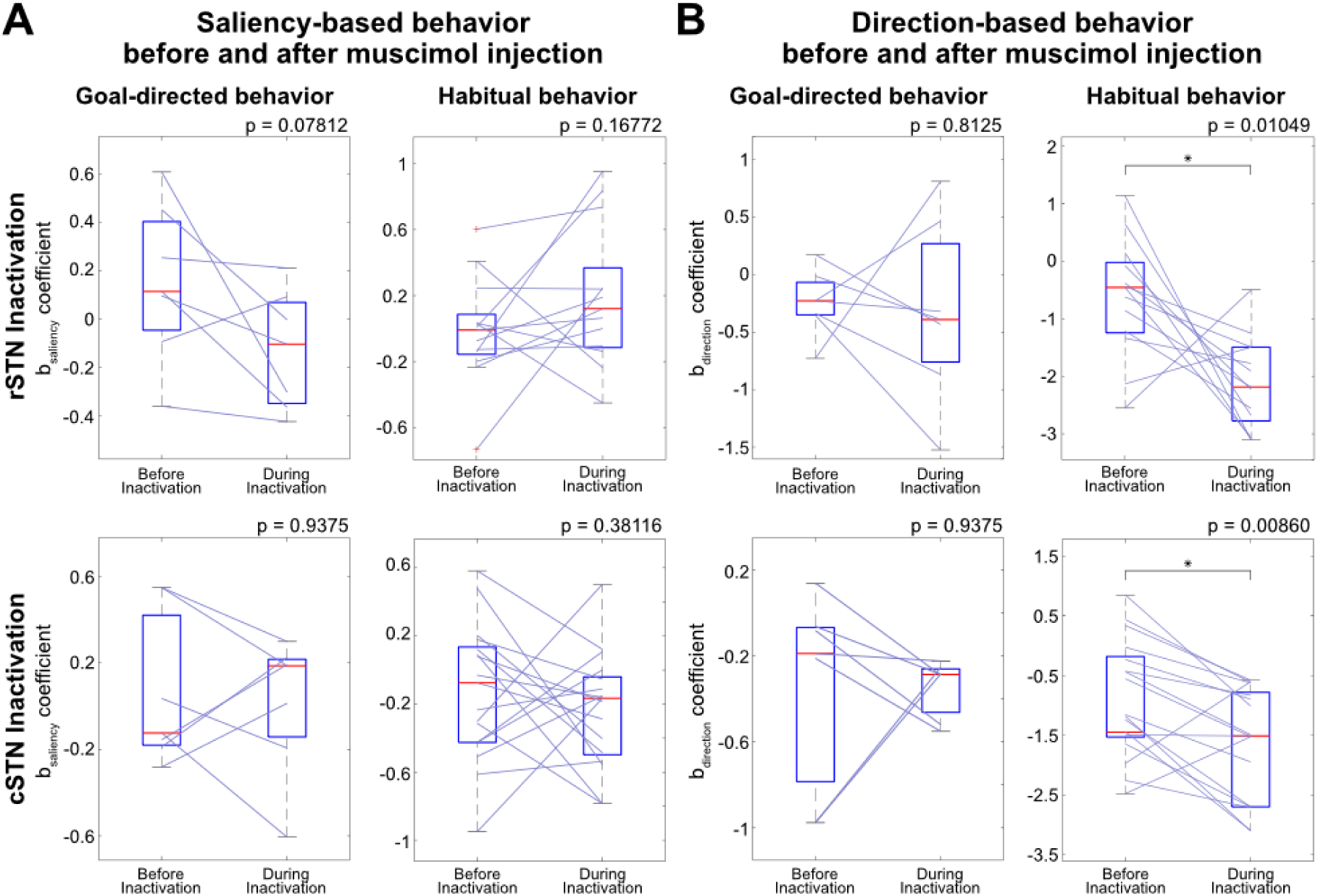
Alterations in direction- and saliency-related behavioral responses following rSTN and cSTN inactivation. (A) Inactivation effect on saliency-based goal-directed (left panels) and habitual behavior (right panels), and (B) inactivation effect on direction-based goal-directed (left panels) and habitual behavior (right panels) following rSTN inactivation (upper panels) and cSTN inactivation (lower panels). Each line indicates the pair of b_saliency_ coefficients (A) and b_direction_ coefficients (B) obtained from logistic regression before and during inactivation. The red line in the blue box represents the mean, and the blue box corresponds to the distribution from the 25th to 75th percentiles. *p < 0.05.

## Notes

### Competing Interest Statement

The authors have declared no competing interest.

